# Environmental nucleic acids: a field-based comparison for monitoring freshwater habitats using eDNA and eRNA

**DOI:** 10.1101/2021.12.01.469845

**Authors:** Joanne E. Littlefair, Michael D. Rennie, Melania E. Cristescu

## Abstract

Nucleic acids released by organisms and isolated from environmental substrates are increasingly being used for molecular biomonitoring. While environmental DNA (eDNA) has received attention recently, the potential of environmental RNA as a biomonitoring tool remains less explored. Several recent studies using paired DNA and RNA metabarcoding of bulk samples suggest that RNA might better reflect “metabolically active” parts of the community. However, such studies mainly capture organismal eDNA and eRNA. For larger eukaryotes, isolation of extra-organismal RNA will be important, but viability needs to be examined in a field-based setting. In this study we evaluate (a) whether extra-organismal eRNA release from macroeukaryotes can be detected given its supposedly rapid degradation, and (b) if the same field collection methods for eDNA can be applied to eRNA. We collected eDNA and eRNA from water in lakes where fish community composition is well documented, enabling a comparison between the two nucleic acids in two different seasons with monitoring using conventional methods. We found that eRNA is released from macroeukaryotes and can be filtered from water and metabarcoded in a similar manner as eDNA to reliably provide species composition information. eRNA had a small but significantly greater true positive rate than eDNA, indicating that it correctly detects more species known to exist in the lakes. Given relatively small differences between the two molecules in describing fish community composition, we conclude that if eRNA provides significant advantages in terms of lability, it is a strong candidate to add to the suite of molecular monitoring tools.

## Introduction

Environmental nucleic acids (eNAs) such as environmental DNA (eDNA) and environmental RNA (eRNA) are emerging as reliable methods for monitoring aquatic biodiversity (Cristescu & Hebert, 2018; Deiner et al., 2017). One concern with the recovery of eDNA revolves around the dynamics of stability and persistence of nucleic acids in the environment that can lead to false detections of local species. There are several possible scenarios through which false positives might result from eDNA; for example, the transport of eDNA molecules from an upstream to a downstream location, or the resuspension of eDNA from a sediment layer which originates from older communities (Corinaldesi, Beolchini, & Dell’Anno, 2008) to the water column. Situations in which it might be difficult to distinguish transported/residual eDNA signal from true signal include detections at low abundance from rare species and at invasion fronts (Jerde, Mahon, Chadderton, & Lodge, 2011). The degree of transported signal relies on the complex interplay among abiotic factors influencing the release, degradation, and persistence of eNAs, the speed and volume of substrate flow, and biotic factors such as biomass, metabolic rates, and behaviour which determine the volume of eNAs released from animal populations. For this reason, it is difficult to predict the extent of transported eNA in each individual scenario. Laboratory and field studies based on eDNA detect signal across distances from several metres to tens of kilometres (Deiner & Altermatt, 2014; Deiner, Fronhofer, Mächler, & Altermatt, 2016; Jane et al., 2015; Jerde et al., 2016; Shogren et al., 2017). This key property of eDNA sampling has important consequences for separating the presence of active communities with molecular monitoring from residual signal.

Recently, it has been proposed that eRNA could be more labile than eDNA and is therefore a candidate molecule for reducing problems associated with transported signal (Cristescu, 2019). The increased lability of eRNA when compared with eDNA is thought to originate from its single-stranded structure, the presence of additional hydroxyl groups allowing for base catalysed hydrolysis (Y. Li & Breaker, 1999), and the ubiquitous presence of exogenous and endogenous RNases (Tan & Yiap, 2009). These characteristics are thought to lead to a faster rate of degradation of eRNA when compared with eDNA; for example, eRNA has a 4-5 hour faster half-life when compared with eDNA (Marshall, Vanderploeg, & Chaganti, 2021). Thus, it may be possible for eRNA signal to distinguish biologically active communities from dead/dormant ones, and local communities from transported molecular signal when eRNA is applied to species monitoring (Barnes & Turner, 2016; Deiner et al., 2017; Pawlowski et al., 2018). For example, eDNA in ballast water was found to contain 57% OTUs assigned to fungi which are thought to represent legacy OTUs, whereas OTUs detected by eRNA included mainly active metazoa and ciliates. Pawlowski et al. (2014) found that DNA recovered greater benthic taxonomic richness when compared with RNA, which could be explained by the detection of previous benthic successions of DNA, as opposed to solely cellularly active taxa (see also Guardiola et al., 2016 for similar findings). The same study found that RNA detected benthic community responses to fish farming to a greater degree than DNA (Pawlowski et al., 2014). Similarly, Dowle et al (2015) found moderately stronger correlations between bacterial RNA OTU data and environmental indices, when compared with DNA OTUs. Although these studies are suggestive, eRNA has received much less attention than eDNA to date, particularly with respect to molecular monitoring of macro-eukaryotes.

Before eRNA is used more commonly in biomonitoring applications, its efficiency of recovery and accuracy of representation of known biological communities must be assessed. Although studies based on organismal RNA are valuable, it is important to examine the viability of extra-organismal RNA recovery under natural field conditions, given that factors such as temperature, acidity, and microbial activity have already been shown to influence eDNA persistence (Sansom & Sassoubre, 2017; Seymour et al., 2018; Strickler, Fremier, & Goldberg, 2015; Tsuji, Ushio, Sakurai, Minamoto, & Yamanaka, 2017). There are many practical considerations when applying molecular monitoring methods in the field, such as the time between sample collection and storage, and choice of buffers used to preserve the molecules until lab work can be undertaken (Deiner, Walser, Mächler, & Altermatt, 2015; Dickie et al., 2018). If eRNA cannot be used in a “field” scenario in the same way that eDNA is, any theoretical benefits due to increased molecule lability would be outweighed by practical disadvantages. Moreover, eRNA may be too labile to be reliably sampled in the field in the same way that eDNA is. The few studies that provide comparisons of community recovery with DNA and RNA metabarcoding focussed on metabarcoding of bulk samples (organismal RNA) which primarily includes smaller eukaryotes and bacteria in marine sediment samples (Brandt et al., 2020; Guardiola et al., 2016; Laroche et al., 2018, 2017; Orsi, Biddle, & Edgcomb, 2013; Pawlowski et al., 2016). Larger animals such as vertebrates (e.g. fish and mammals) might be detected by considering extra-organismal molecules isolated from the environment which are not the primary focus of bulk sample studies. Few studies have specifically considered extra-organismal eRNA, although eRNA was compared to eDNA using ddPCR from two species in laboratory aquaria (Wood et al., 2020), and Miyata et al., (2021) recently used eRNA metabarcoding to detect fishes from two sites in the Naka river, Japan.

In our study, we sampled and filtered water from multiple freshwater lakes, extracting eDNA and eRNA from paired filter halves to compare the community composition of fishes recovered by each molecule. We targeted extra-organismal nucleic acids (eDNA and eRNA) from fish by generating amplicon libraries using the MiFish-U 12S rRNA fragment (Miya et al., 2015). We compared the sampling effort required, species detection success and recovered community composition of these two molecules, and compared both against long-term biomonitoring data collected by conventional methods at these same sites. Sampling was conducted in the summer and again in autumn to capture distinct differences in lake thermal structure (stratification during summer and turnover during autumn). We expected the recovered molecular-based community composition to vary between these two sampling periods, due to differences in water stratification and mixing regimes which control how eDNA is distributed in the lakes, and according to the changing seasonal depth preferences of the fishes (Littlefair, Hrenchuk, Blanchfield, Rennie, & Cristescu, 2021).

## Methods

### Field collection

Samples of eDNA and eRNA were collected in parallel to maximize comparability. We sampled a total of seven lakes (two in 2017 and five in 2018; Table S1) at the IISD Experimental Lakes Area (IISD-ELA), a freshwater research facility in northwest Ontario, Canada. Biomonitoring has been ongoing at IISD-ELA since the 1960s, providing a well-developed knowledge of species composition in this area and in these lakes specifically (Table S2).

Six 500ml water samples were collected from each lake along a depth gradient at the deepest point of each lake, by dividing the water column into six even parts and using an electrical pump and PVC tubing secured to a weight to collect the water at the relevant depth. Collecting on a depth gradient maximises the chance of recovering cold-water fish species such as lake trout (*Salvelinus namaycush*) (Littlefair et al., 2021). Water samples were pumped into sterile Whirl-Pak bags and sealed within an individual large Ziplock bag. Tubing was cleaned between each depth point by pumping one litre of 30% bleach, one litre of distilled water, followed by lake water at the new sampling depth for a two-minute period prior to sample collection. Separate tubing was used for each lake. All samples were immediately transported to the field laboratory in a cooler with ice packs and stored at 4 °C until filtration. Filtering of samples took place within 2-5 hours of sample collection. Water samples were filtered onto 47 mm 0.7 μm pore GF/F filters using an electric vacuum pump and filtering manifold (Pall Corporation, ON, Canada) in a room which was not used for animal work. Each filter was cut into two using an individual pair of forceps and scissors. Half of the filter was immediately frozen at −20°C for DNA analysis and half was preserved in 370μl RLT buffer (Qiagen) with 1% β-mercaptoethanol and then frozen at −20°C for RNA analysis. One negative control of 500 ml distilled water was stored in the cooler and filtered in the same way as the field samples for each lake. In total, 84 eDNA and 84 eRNA samples were taken across the entire study. Filters were shipped on dry ice to McGill University, Montréal for molecular analysis.

### Molecular analysis

Extractions of eRNA were performed from the first half of the filter using the Qiagen RNEasy Mini kit with some modifications to accommodate the filter. Filters were vortexed for 20 seconds and centrifuged in the RLT/ β-mercaptoethanol buffer for 3 minutes at 14,000rpm. A total of 325μl of this buffer was mixed with 325μl ethanol and the rest of the procedure followed the kit protocol intended for extracting total RNA from animal cells. The eRNA was resuspended in two elutions of 30μl RNAse free water to give a final volume of 60μl.

Extractions of eDNA from the second half of the filter were performed using the Qiagen Blood and Tissue kit. We followed the manufacturer’s instructions except that 370μl of ATL buffer was used in an initial overnight incubation step. After extraction, both eDNA and eRNA were preserved at −80°C.

To avoid DNA contamination in the eRNA samples, DNA was digested from 20μl eRNA extracts with the DNA-free™ DNA Removal Kit (ThermoFisher Scientific) following the manufacturer’s instructions and using 2μl DNase I Buffer, 1μl rDNase I and 2μl DNase Inactivation reagent. Samples were checked for residual contaminating DNA using PCR amplification using the MiFish-U primers tagged with Illumina sequencing adapters (Miya et al., 2015). These primers target a hypervariable region of the 12S rRNA locus (163-185bp in length) which has previously been used to characterise the fish community in this area and provides good species level discrimination of ASVs (Littlefair et al., 2021). We used the following PCR chemistry: 7.4μl nuclease free water (Qiagen), 1.25μl 10X buffer (Genscript), 1 mM MgCl2 (ThermoFisher Scientific), 0.2mM GeneDirex dNTPs, 0.05mg bovine serum albumen (ThermoFisher Scientific), 0.25mM each primer, 1U taq (Genscript) and 2μl DNA in a final volume of 12.5μl. We followed a touchdown thermocycling protocol which we have found reduces the amount of non-specific amplification (bacterial taxa) at this locus: 95°C for 3 minutes, 12 cycles of touchdown PCR (98°C for 20 seconds, 66°C for 15 seconds decreasing by 0.2°C each time, 72°C for 15 seconds) followed by 28 cycles with an annealing temperature of 64°C, 72°C extension for 5 minutes. No residual contaminating DNA was found in eRNA samples. A total of 10μl of sample was therefore reverse transcribed into cDNA using the High-Capacity cDNA Reverse Transcription kit (ThermoFisher Scientific) in 20μl reactions following the kit instructions.

We then amplified the cDNA and eDNA in triplicate 12.5μl reactions following the MiFish-U PCR protocol laid out above, and checked amplification using 1% agarose gels with SYBR Safe. We then combined the triplicate reactions into one sample and performed a cleanup with AMPure beads. Cleaned amplicons were then dual-indexed using the Nextera v2 DNA indexes, cleaned again, equimolarised to 3ng/μl and sent for sequencing at Génome Québec, Montréal. Samples were sequenced using 2 x 250bp paired end sequencing with an Illumina MiSeq.

To prevent contamination, we processed the samples in a clean, pre-PCR dedicated lab. Before beginning any eRNA work we thoroughly cleaned benches with 10% bleach solution and RNAse wiper. Laboratory equipment was cleaned with 70% ethanol and RNAse wiper. Negative controls were included at each major step: field sampling, RNA/DNA extraction, reverse transcription, and PCR amplification.

### Bioinformatics

We used custom scripts to remove adapters, merge paired sequences, check quality and generate amplicon sequencing variants (ASVs). Samples were received as demultiplexed fastq files from Génome Québec. Non-biological nucleotides were removed (primers, indices and adapters) using cutadapt (Martin, 2011). Paired reads were merged using PEAR (Zhang, Kobert, Flouri, & Stamatakis, 2014). Quality scores for sequences were analysed with FASTQC (Andrews, 2010). Reads were length filtered between 152-192 bp. Amplicon sequencing variants (ASVs) were generated using the UNOISE3 package (Edgar, 2016), which uses a denoising pipeline to remove sequencing error and to cluster sequences into single variants (100% similarity). The full bioinformatics pipeline is available from https://github.com/CristescuLab/YAAP. After ASVs were generated, we assigned taxonomy using BLAST+ (Camacho et al., 2009) and BASTA (Kahlke & Ralph, 2019), a last common ancestor algorithm. We used a custom reference database which contained only fish known to exist in the Lake of the Woods region (Ontario, CA), downloaded from the NCBI database on 12 August 2018. We also compared our assignments against the full NCBI database and found only one additional fish ASV with the larger database. Other taxonomic groups appeared at very low frequencies when our ASVs were matched against the NCBI database, such as bacterial, mammalian and bird taxa.

### Statistical analysis

All analyses were conducted in R v4.0.2.

#### Bioinformatic filtering

Differences in the final library sizes of eDNA and eRNA filter halves after bioinformatic filtering were analysed by performing a paired *t*-test. We also explored the correlation between library sizes in paired filter halves with a Spearman’s rank correlation test.

#### Sampling effort

For these analyses, ASV count data was converted to incidence data. The number of water samples required to adequately sample species richness was assessed by creating sample accumulation curves using the function specaccum in the “vegan” package (Oksanen et al., 2019). Dataframes were filtered to only include fish species (i.e. the small amount of non-target taxa were removed) in order to draw comparisons with conventional fishing techniques for surveying biodiversity. The differences between eDNA and eRNA accumulation curves were assessed by plotting separate curves for each molecule within each lake and season. Species richness according to conventional techniques was plotted on the graphs with a grey dashed line. We also plotted species accumulation curves with the full dataset (i.e. with non-target taxa included); these are presented in the supplementary material. The ability of each molecule to achieve adequate sampling of species richness was determined by calculating observed sample coverage for each molecule, season and lake combination using iNEXT. We also looked at how species richness varied with increasing sample coverage for each molecule, season and lake combination.

#### Species detection

The fish species composition of the lakes is well known as a result of decades of ongoing monitoring. It was therefore possible to assess the relative performance of eDNA and eRNA to determine species composition against conventional techniques. Fish species were recorded as being present according to conventional techniques if they were consistently detected by typical collection methods (trap netting and short-set gill netting), as part of an ongoing broad scale monitoring program (using sampling procedures as outlined in Rennie et al., 2019, which are typical of sampling efforts in the IISD-ELA lakes in the current study). Using these techniques, we recorded 104 detections with conventional techniques across all the lakes and species. A species was recorded as being present in a lake according to molecular methods if it was detected in at least one of the six water samples taken from that lake. The number of true positives detected by eDNA and eRNA was expressed as a fraction of the total number of conventional detections possible across all lakes and seasons.

We then calculated the true positive rate as the number of detections made with molecular methods as a proportion of true positives and false negatives (which we defined according to the results of conventional sampling). We additionally calculated the false discovery rate as the number of false positives (i.e. species we know are not present in the lakes according to conventional sampling) as a proportion of true positives and false positives (i.e. all detections). True positive and false discovery rates are positive numbers on the scale of 0 – 1, with a higher number indicating a larger proportion of true positives or false discovery in the data. We then used a mixed effects model (fitted with glmmTMB) to examine whether the true positive rate and the false discovery rates differed significantly between eDNA and eRNA, using sampling season as a covariate and fitting the lake as a random effect. We tested all models for overdispersion and examined model residuals using Dharma (Hartig, 2021), and tested the significance of each explanatory term by fitting nested models using the “drop1” command with a chi-squared distribution.

#### Community composition

nMDS was applied to a Bray-Curtis dissimilarity matrix to visually explore the differences in community composition between eDNA and eRNA for each lake. 95% confidence ellipses were drawn around each season/molecule grouping using the ordiplot and ordiellipse functions in vegan. We used the manyglm function in the mvabund package to fit multispecies GLMs for hypothesis testing (Wang, Naumann, Wright, & Warton, 2012). We tested the effects of environmental nucleic acid type (eDNA/eRNA) and season (August/October) as predictors on the community dataset, as well as the interaction between these two factors. When we tested the interaction term, it was not statistically significant, so we removed it and fitted a new additive model with main effects only. We included library size as a log offset to account for library size variation between samples. We tested both poisson and negative binomial distributions and found that the negative binomial distribution removed patterns in the model residuals, so we retained this distribution for our models. We used the anova.manyglm function to retrieve test statistics using adjusted *p*-values to account for multiple testing (i.e. the detection of multiple species). We accounted for the block design of sampling multiple lakes by restricting permutations to within-lake blocks, by supplying a permutation matrix to the bootID argument in the anova function designed using the permute function in vegan.

## Results

### Bioinformatic filtering

Initial library sizes before bioinformatic filtering for eRNA were on average 6.22% smaller than for eDNA. Similar amounts of sequences were removed for both molecule types during the process of adapter removal, pair merging and final trim of primers (Table 1). However, many more sequences were removed during the length filtering step for eRNA when compared with eDNA (eRNA = 25.4% removed, compared with eDNA = 2.43% removed), indicating that the amplification and sequencing of eRNA resulted in more sequences outside the 152-192bp length filter. Despite significantly smaller average library sizes for eRNA compared with eDNA after bioinformatic filtering (paired *t*-test, *t* = 4.09, df = 97, *p* < 0.001), the denoising steps produced similar amounts of ASVs for both molecules (eDNA = 107, eRNA = 115), and similar percentages of sequences mapped onto these ASVs (eDNA = 98.5%, eRNA = 98.4%). There was a moderate but significant correlation in sequence numbers of the filtered library size from paired eDNA and eRNA extracted from the same water sample (Spearman’s rho = 0.539, p < 0.001).

**Table 1:** Sequences present at each step of bioinformatic filtering.

| Bioinformatics step | Mean ± 95% confidence error (per sample) | |
| --- | --- | --- |
|  | DNA | RNA |
| Initial sequence number | 83,972 ± 11,995 | 78,751 ± 12,075 |
| Initial sequence quality score | 32.5 ± 0.61 | 32.0 ± 0.61 |
| Adapter trim (Cutadapt) | 74,493 ± 11134 | 68,107 ± 11,326 |
| Merge reads (Pear) | 74,450 ± 11131 | 67,169 ± 11,397 |
| Trim of remaining reads (Cutadapt) | 74,444 ± 11130 | 68,018 ± 11,318 |
| Length filter | 72,636 ± 11112 | 50,735 ± 9,932 |
| Denoising pipeline | Mean ± 95% confidence error (all samples) | |
| Total sequences after length filtering | 7,118,369 | 4,971,983 |
| Number of non-chimeric ASVs | 107 | 115 |
| Number of reads mapped onto ASVs | 7,012,122 of 7,118,369<br>= 98.5% | 4,890,593 of 4,971,983<br>= 98.4% |

### Sampling effort

Species accumulation curves were more rapid for eRNA taken in October than for eDNA or eRNA in August in 4/7 lakes, but were otherwise inconsistent among lakes, within or across seasons (Figure 1). Generally only a very small number of water samples (three - four) per lake were needed in order to achieve a plateaued species accumulation curve. Compared with conventional techniques, molecular techniques sometimes under- or over-sampled fish species richness, but generally only by one or two species (see below). When considering the entire dataset (i.e. fish and non-target taxa), the curves showed a higher species richness than expected in the lakes based on conventional sampling (Figure S1), primarily because the MiFish-U marker detects small numbers of taxonomic groups other than fish (e.g. zooplankton, human DNA, birds; see below). Mean sample coverage for eDNA was 0.882 and for eRNA was 0.863 (Table S3). Although we did not sample sufficient lakes to conduct a formal statistical analysis on the differences in sample coverage between eDNA and eRNA, there were no noticeable visual trends (Figure S2).

**Figure 1:**
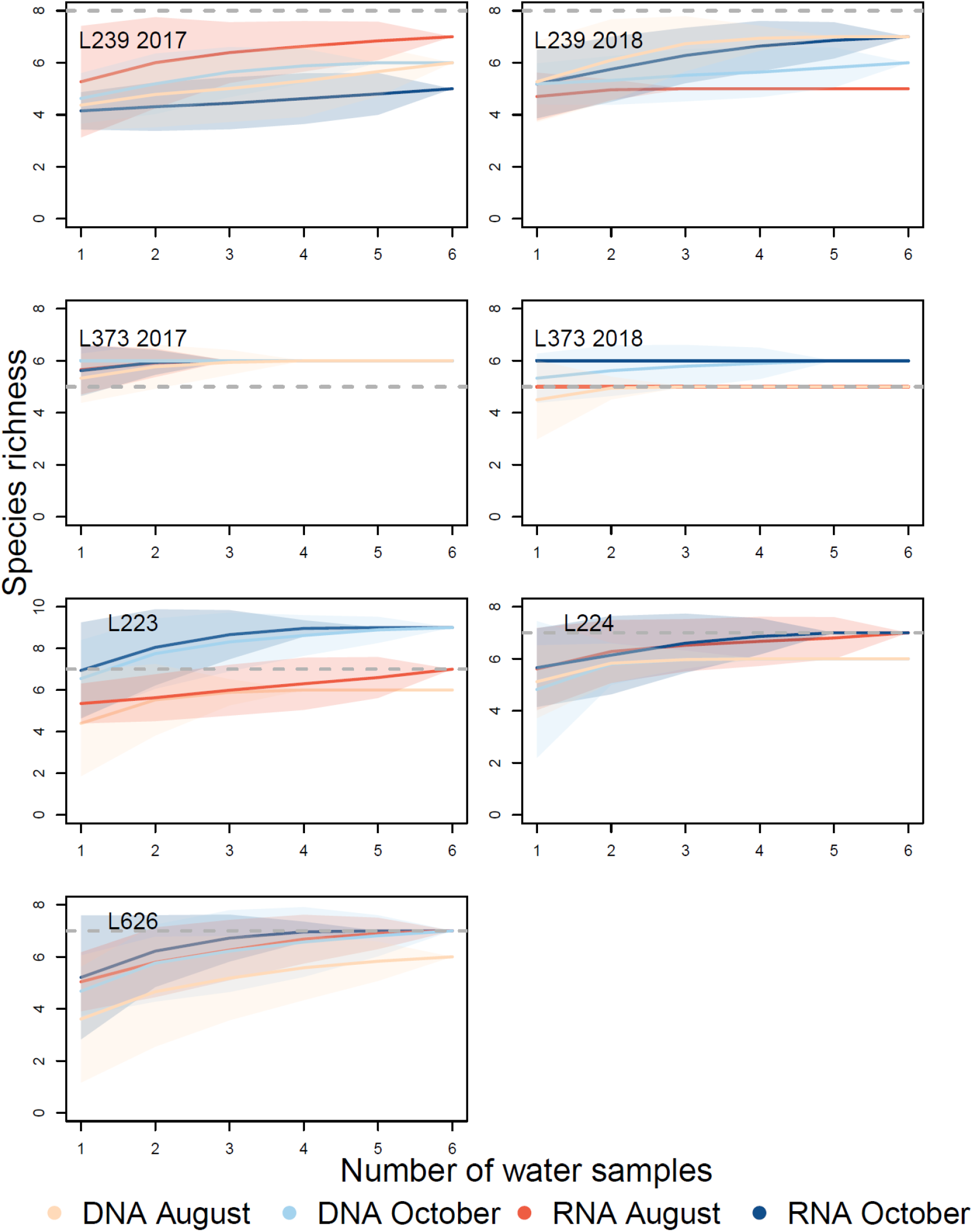
Species accumulation curves for each lake habitat in the study, based on incidence data. The dataframe was filtered to include only fish species present in the northwest Ontario region (i.e. removing non-target ASVs). Separate curves are drawn for ASVs detected with DNA and RNA molecules in the two sampling seasons (August and October), created with random sampling of sites in function specaccum in “vegan”. The grey dashed line indicates the expected number of fish species in that lake according to conventional fishing methods.

### Species detection

The vast majority of sequences (95.8%) in this dataset were assigned to Actinopterygii ASVs. A total of 15 Actinopterygii ASVs were detected by both eDNA and eRNA, with an additional four detected by eRNA only and an additional six detected by eDNA only. Non-target ASVs were also generated, although only a relatively small percentage of sequences in the dataset were actually assigned to these ASVs (4.23%). Non-target ASVs varied between the two molecules: eRNA detected a greater incidence of bacterial, algal and arthropod ASVs, and eDNA detected more mammalian and unassigned ASVs.

When using the database consisting of fish found in the northwest Ontario region, ASVs could generally be assigned at species level. There were two fish species which were not detected by either molecule (*Culaea inconstans* and *Rhinichthys cataractae*). Moreover, *Chrosomus neogaeus* and *Chrosomus eos* could only be detected at genus level from a single ASV: species specific identification is not possible due to the presence of mitochondrial hybrids in the region (Mee & Taylor, 2012). We therefore counted detections only once for the *Chrosomus* genus to avoid double counting.

There were very small amounts of sequences detected in negative controls after bioinformatic filtering. These controls had an average library size of 178, compared with eDNA/eRNA libraries which had an average size of 69,408. Of these, 91.7% of sequences in the negative controls matched fish from northwest Ontario (rather than other taxa such as bacteria or mammals). In almost all cases, the sequences matched the species composition from the lake that the negative control originated from, indicating that contamination of the negative control originated from within the lake sampled, rather than across-lake contamination or tag jumping. We did not find any amplification in the eRNA after the use of the DNA removal kit before conversion to cDNA.

Sampled eRNA had a small but significantly greater true positive rate than eDNA, indicating that eRNA correctly detected more of the species known to exist in the lakes based on conventional sampling (Figure 2A, eRNA true positive rate: 0.692 per sample, eDNA true positive rate: 0.648 per sample, *p* = 0.0043). There was no difference in false discovery rate between the two molecules; i.e., neither molecule detected more false positives as a proportion of all detections (Figure 2B, eRNA: 0.052, eDNA: 0.046, *p* = 0.568). There was a significantly lower false discovery rate in August samples when compared with those collected in October (August: 0.030, October: 0.069, *p* = 0.0004). Usually false positive detections were of a low read count (Table S2).

**Figure 2:**
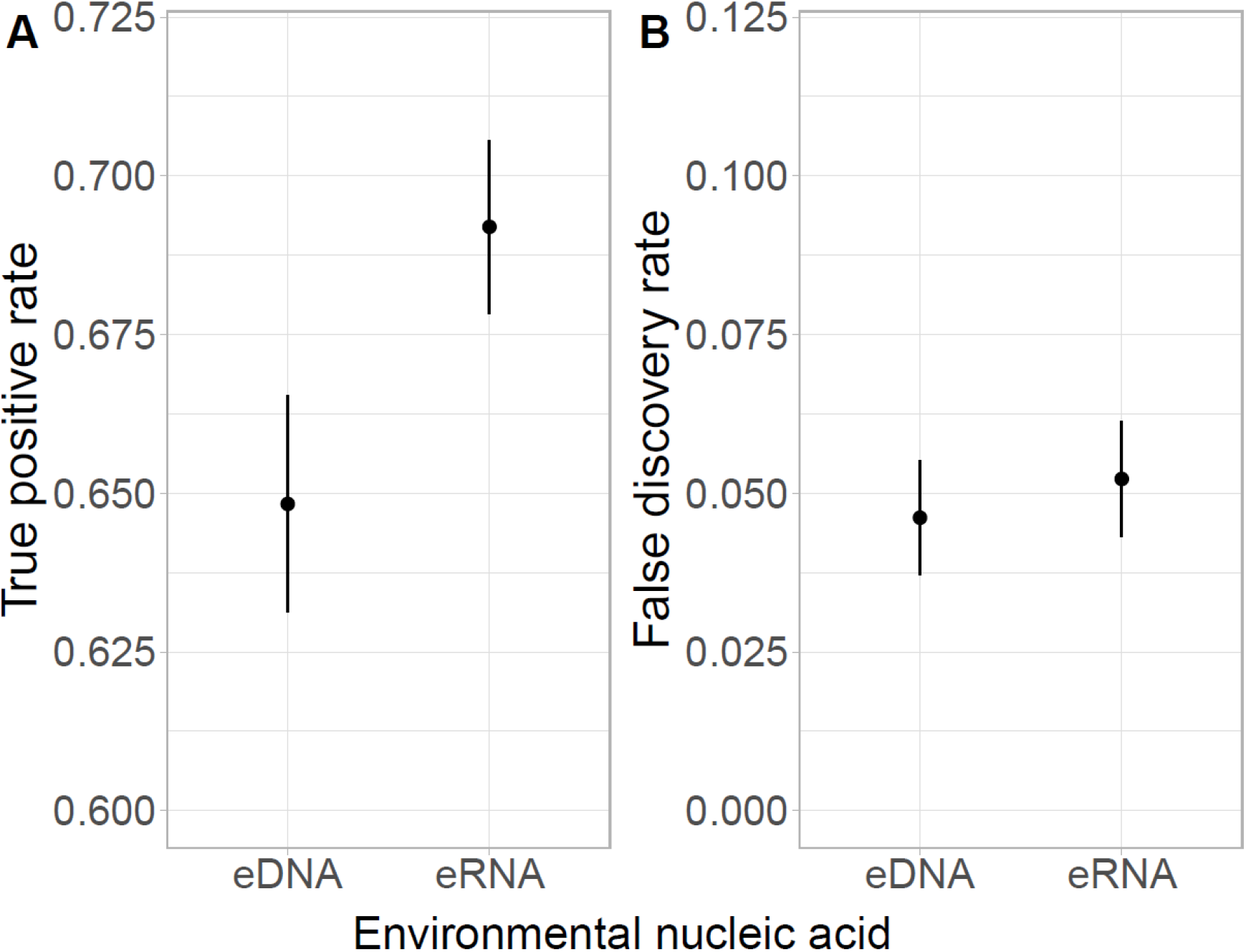
A) True positive rate and B) false discovery rate as measured by eDNA and eRNA. eRNA had a significantly higher true positive rate than eDNA. True positive rate is defined as the proportion of true positives out of true positives and false negatives combined. There was no significant difference in false discovery rate between eDNA and eRNA. False discovery rate is defined as the proportion of false positives out of all detections (i.e. true positives and false positives together). Error bars are standard errors of the mean.

### Community composition

Using NMDS as a visual technique to explore the differences in community composition as explained by different predictors, the largest differences in community composition were generated by the differences in species composition between lakes (Figure S3). Samples collected in October were more similar to each other in terms of community composition than samples collected in August (Figure S4), although these two distributions were nested within each other. In the overall dataset, eDNA and eRNA samples detected largely similar communities, although eRNA samples were slightly more similar to each other than eDNA samples (Figure S5). Within a single lake, there was variation in whether eDNA and eRNA detected similar community compositions (Figure 3). Although consistent differences between the two molecules were not evident, there did seem to be a stronger seasonal effect, as samples that were collected in August were often more dissimilar to each other than samples collected in October. This is reflected in a larger and partially non-overlapping 95% confidence ellipse for these groups of samples.

**Figure 3:**
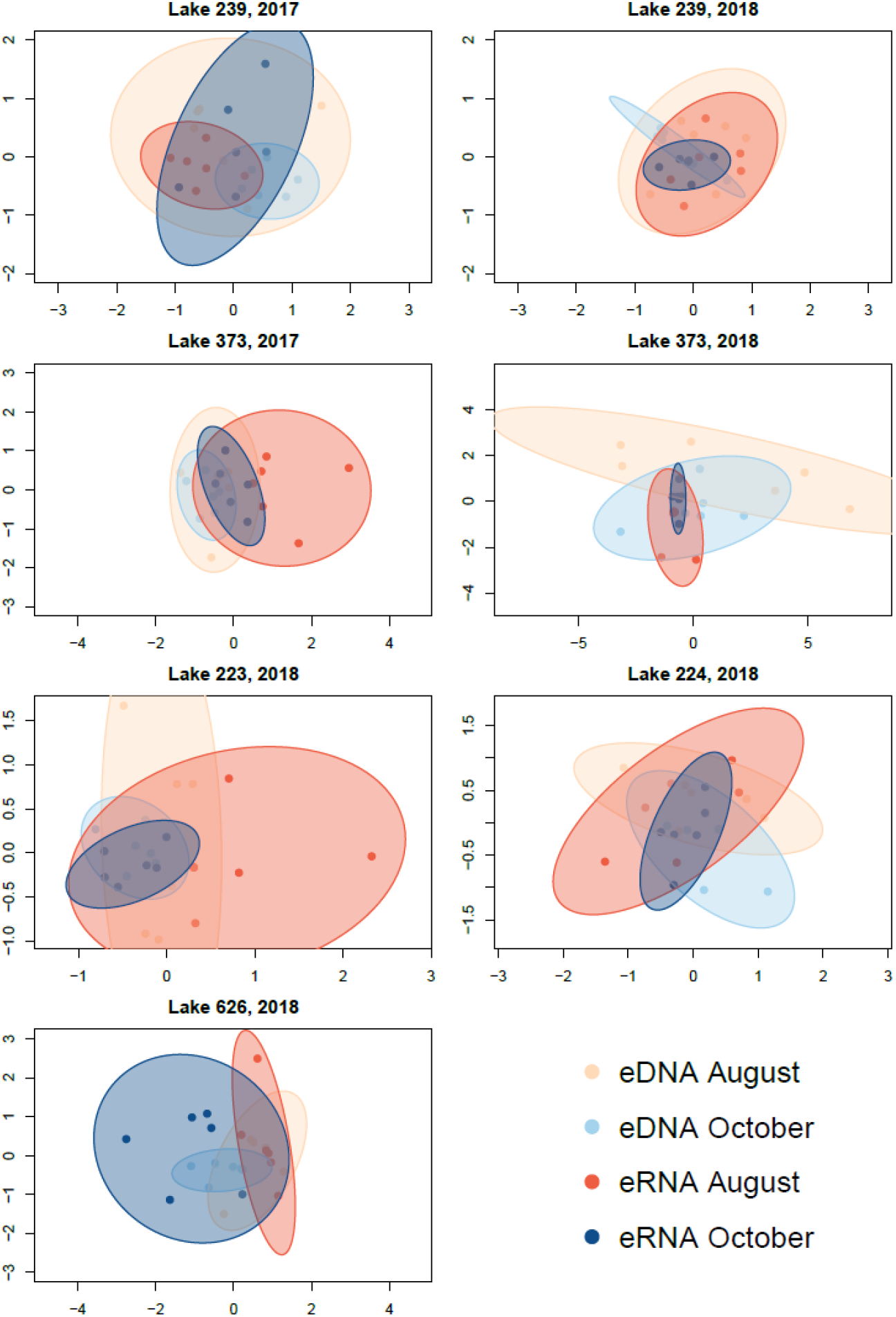
nMDS plots for each lake showing community dissimilarities for both eDNA and eRNA in August and October. New nMDS are run for each lake; thus, scaling varies for each image. Ellipses are 95% confidence intervals coloured according to environmental nucleic acid and season.

We visually explored the proportion of sequences in each eDNA and eRNA sample which belonged to each species or taxa in August (Figure 4A) and October (Figure 4B). We calculated the proportion of sequences belonging to each species out of the total library size per filter. In some taxa, eDNA and eRNA samples had strikingly similar proportions of sequences within each library, for example white sucker (*Catostomus commersonii*), finescale/northern redbelly dace (*Chrosomus spp.*), slimy sculpin (*Cottus cognatus*), and fathead minnow (*Pimephales promelas*) (see also Table S2). In a few species, there were differences between the proportions of nucleic acids; for example, there was always a higher proportion of *Coregonus artedi* eDNA sequences compared to eRNA, and always a lower proportion of *Perca flavescens* eDNA sequences compared to eRNA sequences. There were also seasonal differences in the proportion of sequences belonging to each species; for example, there were more sequences belonging to lake trout (*Salvelinus namaycush*) in October compared with August, which reflects the spawning times and patterns of habitat occupancy for this fish (Littlefair et al., 2021). Similarly, there were also seasonal effects on sequence numbers of *Coregonus artedi* and *Cottus cognatus* which are both cold water fish (higher concentrations in October), and much lower concentrations of *Perca flavescens* sequences in October. These visual differences were reflected by the results of the multispecies GLM, which retained significant effects for molecule type (df = 1, 166, deviance = 284.7, p = 0.02) and season (df = 1, 165, deviance = 296.4, p = 0.005). There was no significant effect of the interaction between molecule type and season on the numbers of sequences, indicating that eDNA and eRNA detection of taxa did not respond differently between the two seasons.

**Figure 4:**
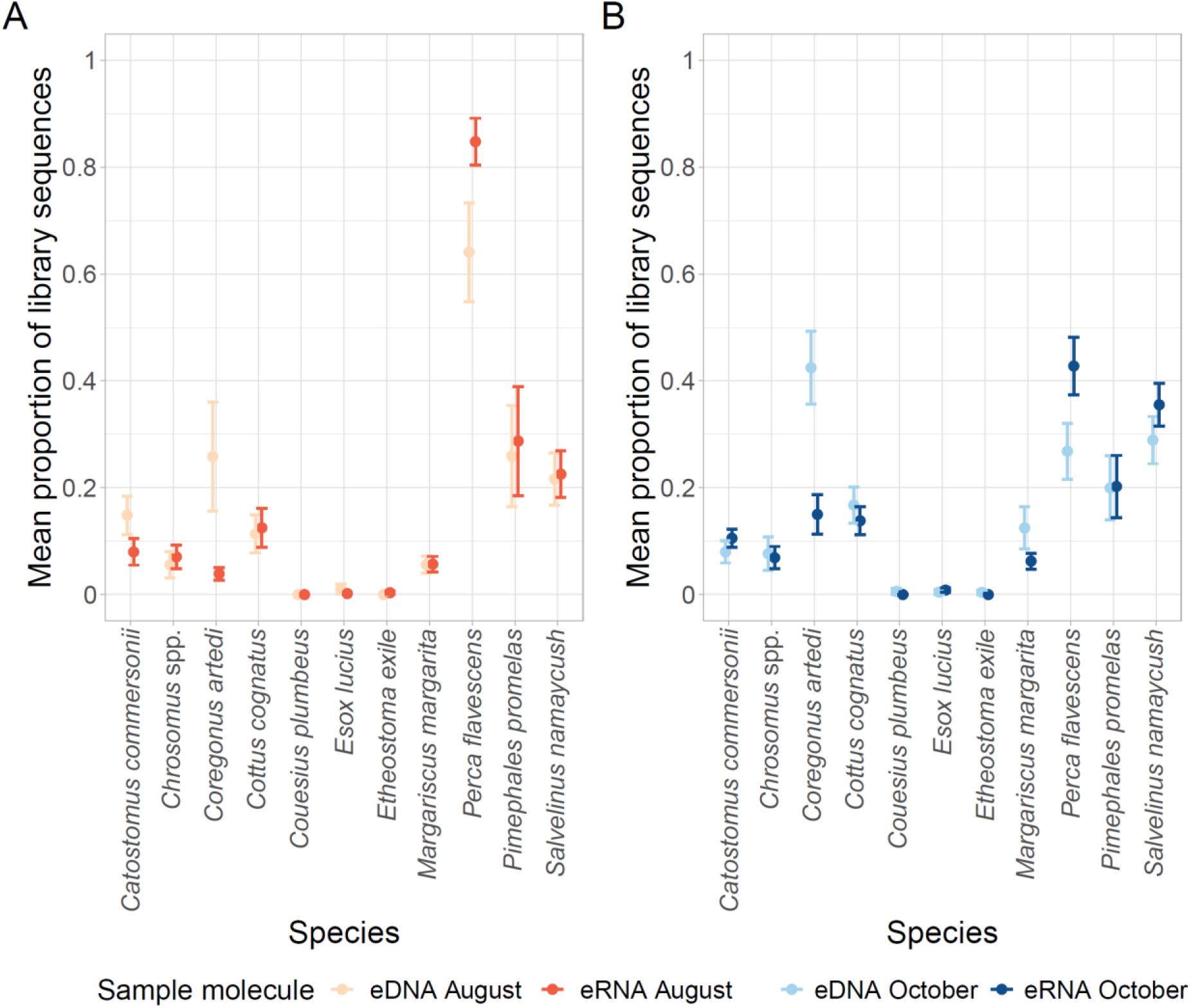
Proportional composition of ASVs per filter for each fish species for A) August and B) October. Where there is more than one ASV per species, these are grouped together under the species name. Error bars are standard errors of the mean.

## Discussion

Traditionally, RNA has been thought of as a very labile molecule, too difficult to extract and preserve in a field setting. However, we have shown here that eRNA achieves similar rates of macroeukaryotic species detection when compared with eDNA within the context of sampling for environmental assessment, and in fact had a slightly higher rate of true positive detection per sample than eDNA. The majority of RNA studies to date have focused on metabarcoding of bulk samples (organismal RNA), but the choice of fish as a study organism shows that it is possible to detect animals based simply on extra-organismal RNA released into the environment. This is the first paper to solely focus on the release of extra-organismal RNA in lakes with comparisons to both eDNA and well-documented conventional monitoring techniques, and these comparisons support its application more broadly to assess species presence/absence and ecosystem functioning.

Extra-organismal RNA is stable enough to collect with comparable field techniques used with extra-organismal DNA. Across the entire study, all target fish species which were detected with eDNA could also be detected with eRNA. Per sample, the true positive rate of detection was significantly higher with eRNA than eDNA, reflecting the results of Miyata et al., (2021), although the difference in detection rate between the two molecules was small. This is consistent with studies based on bulk DNA and RNA which report substantial overlap between OTU or species detections by the two molecules (e.g. Pochon, Zaiko, Fletcher, Laroche, & Wood, 2017). In our study, field collection involved pumping water from the lake, transporting the water samples back to the field laboratory, and filtering them, which took place between two and five hours after collection, a protocol typical of many eDNA studies (Balasingham, Walter, Mandrak, & Heath, 2018; Bylemans, Gleeson, Hardy, & Furlan, 2018; Hänfling et al., 2016; Jeunen et al., 2019; J. Li, Lawson Handley, Read, & Hänfling, 2018; Stat et al., 2018; Zhang et al., 2020). Here, we have shown that extra-organismal eRNA can withstand comparable field methods and can be filtered and sequenced from the water column, performing comparably to eDNA for species detections when compared with conventional monitoring techniques.

While false negatives occurred for both eDNA and eRNA in certain lakes when molecular results were compared to the known species composition of the lake ascertained by conventional techniques, the incidence of these false negatives seemed to be linked to the ecology of these species rather than the type of nucleic acid molecule. For example, fish species which could not be detected in certain lakes were almost always recorded by the biomonitoring program as being at rare or moderate levels of abundance in that particular lake (e.g., *Couesius plumbeus* in lake 626). Moreover, species which favoured a littoral habitat or which live in small inlets around the lakes (e.g., *Chrosomus* spp, *Catostomus commersoni*, and *Esox lucius*) were also recorded as false negatives in some lakes. This is perhaps not surprising given that sampling took place at the centre point of the lakes, well away from the shoreline. The two fish which were reported as being present by conventional techniques but not detected by molecular sampling in any lake habitat (*Culaea inconstans* and *Rhinichthys cataractae*) were likewise recorded as being at “rare” levels of abundance and were also species which favoured littoral or inlet habitats around the periphery of the lakes. For example, only one *Rhinichthys cataractae* was caught in the survey in 2017, and none have been caught in subsequent years.

There was no significant difference in the false discovery rate between eDNA and eRNA. False discovery rate is defined as the number of false positives as a proportion of the sum of true positives and false positives. Some studies have proposed that eRNA might detect cellularly active taxa only, as opposed to dead and dormant taxa or resuspended sedimentary DNA, and thus minimise the false discovery rate when compared with eDNA (Dowle et al., 2015; Pawlowski et al., 2014; Pochon et al., 2017; Visco et al., 2015, although see Brandt et al., 2020), although to date laboratory degradation experiments indicate that eRNA might not degrade significantly faster than eDNA (Wood et al., 2020). Further studies in a field setting will be needed to determine the advantages of eRNA of overcoming false positives detected by eDNA sampling, should they exist. In semi-natural settings, experiments have been performed using caged animals or artificial spikes of DNA to assess the effects of time or distance on the degradation of the DNA signal (Harper, Anucha, Turnbull, Bean, & Leaver, 2018; Jane et al., 2015; Pilliod, Goldberg, Arkle, & Waits, 2014). Observational field studies have also provided evidence that DNA flows downstream from populations (Deiner & Altermatt, 2014; Deiner et al., 2016). These methodologies could be performed in parallel with eRNA to analyse whether the use of eRNA improves the false positives currently detected by eDNA. Some of the lakes that we examined in this study are headwater lakes (Table S1), meaning that there is no opportunity for water and nucleic acids to flow from upstream populations, so the lack of difference in false discovery rate between eDNA and eRNA is perhaps unsurprising in this context.

We did not specifically analyse the degradation time of eRNA in our study, but it is noteworthy that a reliable signal from this nucleic acid could still be detected two to five hours after collection in the field and storage in refrigerated conditions. This and several other lines of evidence point to extra-organismal RNA being more stable than previously thought (Cristescu, 2019; Torti, Lever, & Jørgensen, 2015). It may be that the eRNA is temporarily stable inside cells, organelles or vesicles; a study of long noncoding RNA (lncRNA) found that its half-life within cells ranged from <2 to >16 hours (Clark et al., 2012). Although Marshall et al., (2021) found that eRNA had a 4-5hour faster half-life than eDNA, the half-life was still within 8.84 – 13.54 hours (depending on marker selection and RNA type), which was within the collection window of our study. Alternatively, eRNA could be combined with organic or inorganic particles within the water column which aids with stability as with eDNA; for example, Wood et al., (2020) found that eRNA could still be detected in biofilms from 4/15 aquaria at the end of a 21-day experiment. There have also been suggestions that RNA might preserved for long time periods in sediments by binding to sedimentary particles (Cristescu, 2019; Orsi et al., 2013; Torti et al., 2015). For eRNA to be a successful complement to eDNA to resolve spatial-temporal issues in species detection it must be stable enough to be detected, but ideally degrade faster than eDNA, in order to provide a detection signal which is in closer geographic proximity to the population in question.

We found strong effects of season and molecule type on the proportion of sequences assigned to different fish species in the libraries. In many (but not all) cases, this seems to correspond to aspects of species biology and abundance in the lakes. We found higher proportions of sequences for both molecule types in October for cold water species such as lake herring (*Coregonus artedi*), slimy sculpin (*Cottus cognatus*), and lake trout (*Salvelinus namaycush*) (Figure 4B). The October sampling corresponded to the spawning time of lake trout, an event which has been linked to the creation of eNAs (Tillotson et al., 2018), and although the other two species were not spawning during October, their activity expands from deep, hypolimnetic waters to the entire lake as lake surface temperatures fall below thermal maxima and towards optimum temperatures for these species (Hasnain, Escobar, & Shuter, 2018). Across both seasons, there were high levels of yellow perch (*Perca flavescens*) sequences, with some libraries containing >90% reads assigned to yellow perch. This may reflect the greater abundance of yellow perch in these lakes relative to other species. The numbers of yellow perch sequences in August were particularly high; this may be due to the relative inactivity of other species in warmer parts of the lake during the summer, as the proportion of sequences from one species affects the proportion of sequences from others in the sequencing libraries. There were also consistent effects of molecule type on the proportions of sequences which was largely consistent across seasons; for example, high levels of lake herring eDNA relative to eRNA, and high levels of yellow perch eRNA relative to eDNA in both August and October. However, for many species, the consistency of the relative levels of eDNA and eRNA was surprisingly strong, across both seasons. The reasons for these patterns are, as yet, unknown.

Comparing workflows will be important when assessing the relative strengths of these two molecules. Surprisingly, eRNA was robust to a typical eDNA field protocol, which involved a time lag of 2-5 hours between collection and the filtering and storage of filters (typical of many eDNA studies). Possible increases in yield could be achieved by adapting or applying recent innovations which involve the capture and storage of molecules *in situ* or direct sequencing in the field (Truelove, Andruszkiewicz, & Block, 2019). We applied equivalent levels of care to preserving the stability of both molecules after collection; for example, keeping filters frozen at all times, shipping on dry ice, use of diluted bleach and RNase in the laboratory to clean surfaces and equipment, and working with samples on ice. Inherently, the molecular workflow involves some differences between the two molecules. We chose to use spin column based kits for extractions (Blood and Tissue kit for DNA, RNeasy kit for RNA, both manufactured by Qiagen), as these methods have been shown to yield some of the highest quantities of RNA (Tavares, Alves, Ferreira, & Santos, 2011). Alternative options are provided by kits which co-extract the two molecules together such as the ZR-Duet DNA/RNA MiniPrep kit from Zymo Research (as used in Pochon, Zaiko, Fletcher, Laroche, & Wood, 2017). Sampled eRNA then requires additional steps to convert the molecule into cDNA for sequencing, involving DNA digestion and reverse transcription, before performing an equivalent PCR amplification as in traditional DNA metabarcoding. These additional steps might involve the loss of molecule yield. We found that the final recovered sequence counts were lower for eRNA libraries – given that an equimolar amount was added to the metabarcoding libraries, this loss of molecules possibly occurred through the removal of low-quality reads during the bioinformatics pipeline. As a final note, these additional molecular steps mean that the extraction and processing of eRNA is more costly per sample in terms of kits and personnel time in comparison to eDNA, which may be an important consideration currently when deciding the relative benefits between eDNA and eRNA in field studies.

However, the development of newer sequencing technologies mean that the simplification of this workflow is on the horizon. Future sequencing technologies will mean that RNA can be sequenced directly without conversion to cDNA or with PCR bias which results from amplification steps. The possibility of starting with low-input amounts means that this might be particularly suitable for eRNA applications, and this technology has already been applied to microbial mock communities (Nicholls, Quick, Tang, & Loman, 2019). Some features which will be of interest to eDNA/eRNA scientists are still in development at the time of writing, such as the ability to multiplex samples with direct RNA sequencing kits, but further advances will be anticipated with interest.

## Acknowledgements

We are indebted to many IISD Experimental Lakes Area students and staff for maintaining records of field data and for logistical assistance with this project. S Michaleski, R Henderson, P Bulloch, M Haust, C Jackson and A McLeod contributed specific field assistance to this project, and L Hrenchuk oversaw fieldwork. K Sandilands provided the pumping equipment used in this project. Dr JS Hleap provided bioinformatics support to this project. M Harris provided training on the eRNA extraction and reverse transcription protocols used in this work. This work was funded by a Mitacs Accelerate Industrial Fellowship (JEL), an NSERC Collaborative Research and Development award (MEC), Canada Research Chair and NSERC Discovery awards to MEC and MDR, Québec Centre for Biodiversity Science Excellence award (JEL), the WSP Montréal Environment Department and in-kind support from the IISD Experimental Lakes Area and Fisheries & Oceans Canada. We thank Jean Carreau and Patrick LaFrance of WSP Montréal for useful discussions on the topics of eDNA and biomonitoring.

## Animal permits

Scientific collection permits to facilitate fish collections were authorised through the Ontario Ministry of Natural Resources and Forestry. Handling of fish was carried out under the authority of the Government of Canada through Fisheries and Oceans Canada (prior to and including 2013), and with the approval of Animal Care Committees through the University of Manitoba (2014; permit #F14-007) and Lakehead University (2015–present, permit #s 1464656 and 1466997).

## Data Accessibility statement

Raw fastq files, sample x ASV tables, and the sequence composition of the ASVs are openly available at Dryad (xxx). Scripts to process bioinformatic data are available from https://github.com/CristescuLab/YAAP.

## Author contributions

JEL and MEC designed the research, all authors contributed to funding the research, JEL performed the molecular work and analysis, MDR contributed field data, JEL wrote the first draft of the manuscript and all authors contributed the writing and editing of the manuscript.

## Notes

### Competing Interest Statement

The authors have declared no competing interest.

