## Supporting Information for "Environmental nucleic acids: a field-based comparison for monitoring freshwater habitats using eDNA and eRNA"

Table S1: Physical properties and locations of the study lakes.

| Lake | Coordinates (dd.ddddd) | Size (ha) | Maximum depth (m) | Inflows | Outflows | Sampling months |
|---|---|---|---|---|---|---|
| **239** | N 49.39493<br>W 093.43211 | 56.1 | 30.4 | 3 | 1 | July 2017<br>October 2017<br>August 2018<br>October 2018 |
| **373** | N 49.44422<br>W 093.47589 | 28.0 | 21.5 | Headwater lake | 1 | July 2017<br>October 2017<br>August 2018<br>October 2018 |
| **223** | N 49.698333<br>W 093.70778 | 27.3 | 14.4 | 1 | 1 | August 2018<br>October 2018 |
| **224** | N 49.69000<br>W 093.71694 | 25.9 | 27.4 | 1 | 1 | August 2018<br>October 2018 |
| **626** | N 49.75279<br>W 093.79799 | 25.8 | 13.2 | Headwater lake | 1 | August 2018<br>October 2018 |

Table S2: Fish species composition of the study lakes determined by conventional sampling methods (trap netting, gillnetting, etc) over several decades (represented by X), eDNA (represented by counts in bold text) and eRNA (represented by counts in italic text). Read counts are summed from the six samples taken in August and six taken in October (i.e. both seasons are combined). Note that *Chrosomus neogaeus* and *Chrosomus eos* can be distinguished morphologically but not genetically at these sites, due to the presence of mitochondrial cybrids in this region.

| | **Lake** | | | | | | |
|---|---|---|---|---|---|---|---|
| **Species** | **239 2017** | **239 2018** | **373 2017** | **373 2018** | **223** | **224** | **626** |
| *Couesius plumbeus* | **0**<br>*0* | **0**<br>*0* | **516**<br>*490* | **492**<br>*970* | X<br>**10518**<br>*38* | **0**<br>*0* | X<br>**0**<br>*0* |
| *Pimephales promelas* | **0**<br>*0* | **0**<br>*0* | **0**<br>*0* | **0**<br>*0* | X<br>**176980**<br>*67709* | X<br>**161375**<br>*88969* | **0**<br>*0* |
| *Chrosomus eos* | | | X | X | | X | |
| *Chrosomus neogaeus* | X | X | X | X | X | X | X |
| *Chrosomus (ASV assigned at genus level)* | **0**<br>*0* | **0**<br>*0* | **312261**<br>*219825* | **164183**<br>*146647* | **3413**<br>*2452* | **57**<br>*58452* | **3**<br>*7925* |
| *Catostomus commersonii* | X<br>**5403**<br>*13285* | X<br>**5696**<br>*1212* | X<br>**169247**<br>*122684* | X<br>**226348**<br>*134552* | X<br>**141288**<br>*78771* | X<br>**140630**<br>*74740* | X<br>**12797**<br>*29658* |
| *Etheostoma exile* | X<br>**4534**<br>*3743* | X<br>**10**<br>*3* | **0**<br>*0* | **0**<br>*0* | **0**<br>*0* | **0**<br>*0* | **0**<br>*0* |
| *Culaea inconstans* | **0**<br>*0* | **0**<br>*0* | **0**<br>*0* | **0**<br>*0* | X<br>**0**<br>*0* | X<br>**0**<br>*0* | **0**<br>*0* |
| *Esox lucius* | X<br>**1**<br>*1312* | X<br>**16282**<br>*7850* | **0**<br>*0* | **0**<br>*0* | **0**<br>*0* | **0**<br>*0* | **0**<br>*0* |
| *Perca flavescens* | X<br>**507875**<br>*659011* | X<br>**323232**<br>*401937* | **0**<br>*0* | **0**<br>*0* | **11**<br>*85* | **0**<br>*4* | X<br>**444645**<br>*377868* |
| *Margariscus margarita* | **0**<br>*0* | **0**<br>*0* | X<br>**169519**<br>*84264* | X<br>**117510**<br>*117259* | X<br>**88076**<br>*10936* | X<br>**8830**<br>*1264* | X<br>**13383**<br>*5727* |

*Continued*

Table S2 continued

| | Lake | | | | | | |
|---|---|---|---|---|---|---|---|
| **Species** | **239 2017** | **239 2018** | **373 2017** | **373 2018** | **223** | **224** | **626** |
| *Rhinichthys cataractae* | **0** *0* | **0** *0* | **0** *0* | **0** *0* | **0** *0* | **0** *0* | X* **0** *0* |
| *Coregonus artedi* | X **209182** *65688* | X **531102** *95526* | **0** *0* | **0** *0* | **1** *15* | **0** *0* | **14** *33* |
| *Salvelinus namaycush* | X **38305** *113117* | X **153563** *87681* | X **79406** *225605* | X **466784** *397582* | X **476637** *88440* | X **99206** *195634* | X **484842** *160259* |
| *Cottus cognatus* | X **162963** *66775* | X **183553** *133882* | X **31** *106224* | X **80841** *33835* | X **211308** *215997* | X **199646** *57306* | X **39** *1062* |

* *Rhinichthys cataractae* are currently very rare compared to historic densities in lake 626; only one was caught in 2017, none in 2018 or 2019 or since (vs. dozens in 2014 and 2015).

Table S3: Sample coverages for each environmental nucleic acid and season, per lake. All sample coverages were calculated using the iNEXT package in R.

| | **eDNA - Aug** | **eDNA - Oct** | **eRNA - Aug** | **eRNA - Oct** |
|---|---|---|---|---|
| Lake 239 2017 | 0.8803 | 0.8780 | 0.8727 | 0.9580 |
| Lake 239 2018 | 0.8803 | 0.9173 | 0.8186 | 0.8663 |
| Lake 373 2017 | 0.9179 | 0.9909 | 0.8310 | 0.9110 |
| Lake 373 2018 | 0.6867 | 0.8126 | 1.0000 | 0.8735 |
| Lake 223 | 0.9194 | 0.9414 | 0.8722 | 0.8941 |
| Lake 224 | 0.8227 | 0.8262 | 0.7052 | 0.8889 |
| Lake 626 | 0.9745 | 0.9020 | 0.8898 | 0.7048 |

Figure S1: Species accumulation curves for each lake habitat in the study, based on incidence data. The dataframe includes both target and non-target ASVs. Separate curves are drawn for ASVs detected with DNA and RNA molecules in the two sampling seasons (August and October), created with random sampling of sites in function specaccum in “vegan”.

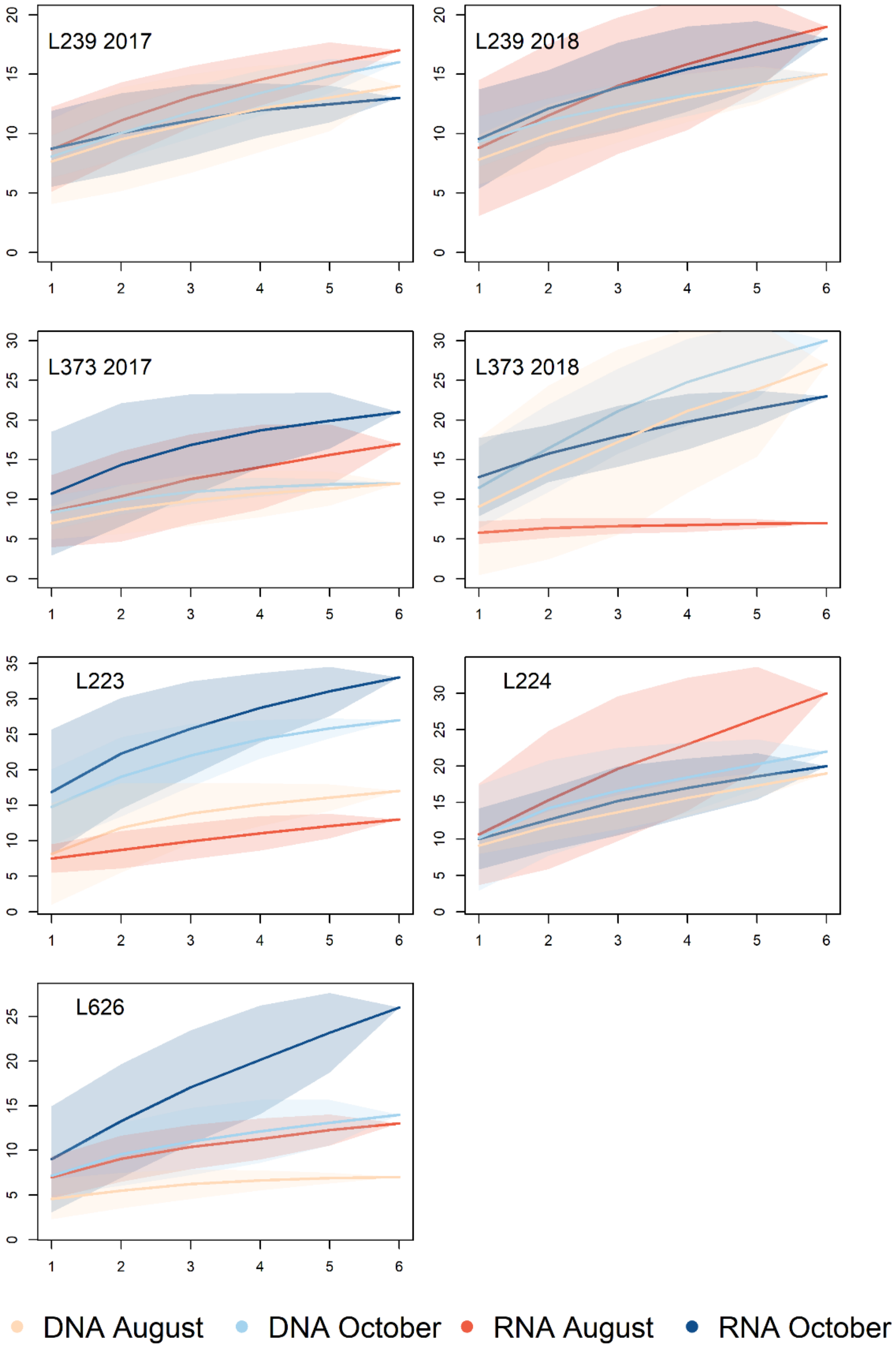

Figure S2: Box plot demonstrating sample coverage did not vary according to the environmental nucleic acid (eDNA or eRNA) or the season the samples were collected in.

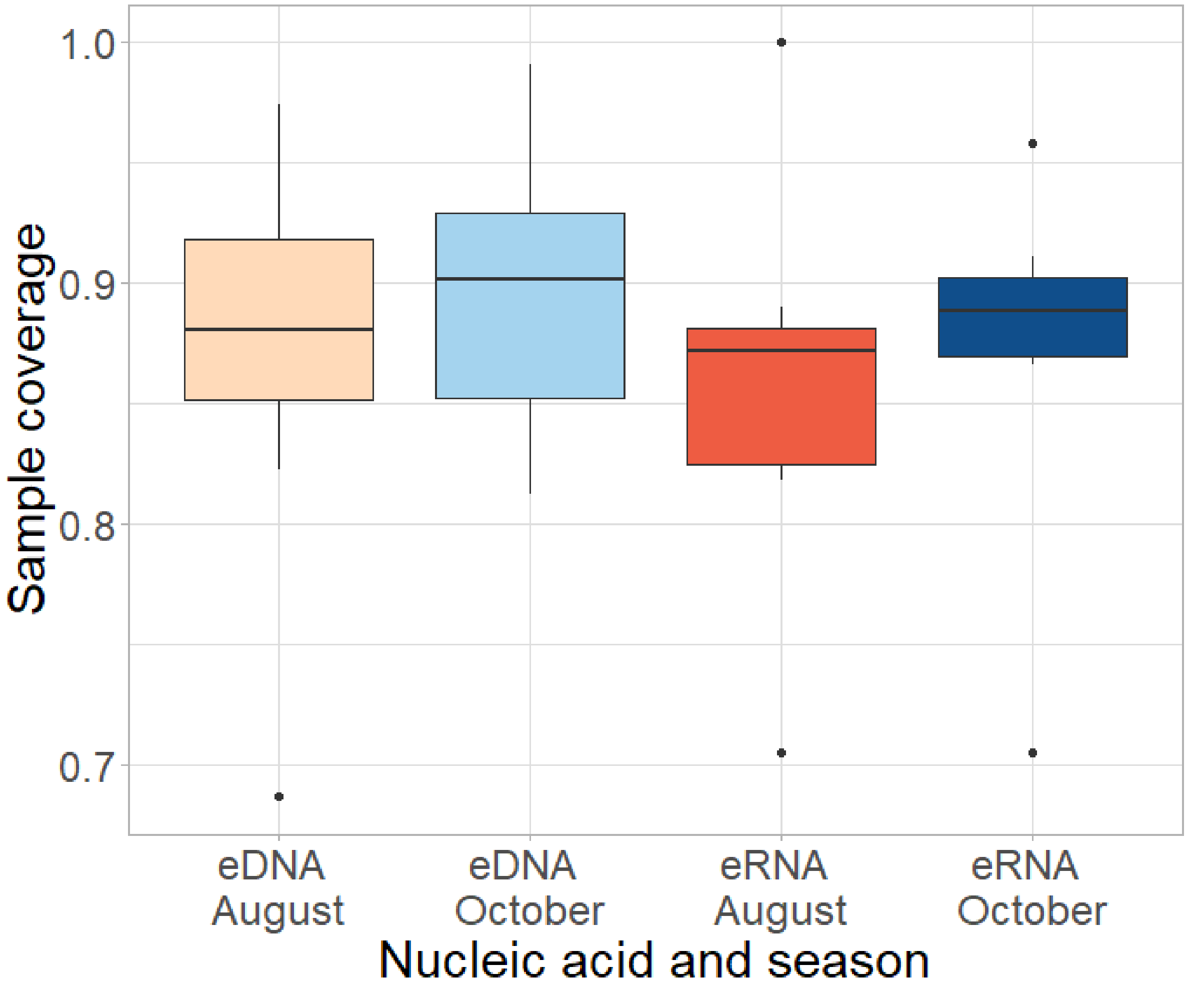

Figure S3: nMDS was performed on the entire dataset (eDNA and eRNA samples together) to visualise the principle drivers of sample similarity. The main differences in the dataset were driven by the different lake habitats, most likely due to the different fish species composition of the lakes (95% confidence ellipses coloured for each lake). The nMDS 1 axis shows separation of lakes according to the presence or absence of yellow perch (*Perca flavescens*). The nMDS 2 axis shows separation of lakes according to the presence or absence of fathead minnows (*Pimephales promelas*).

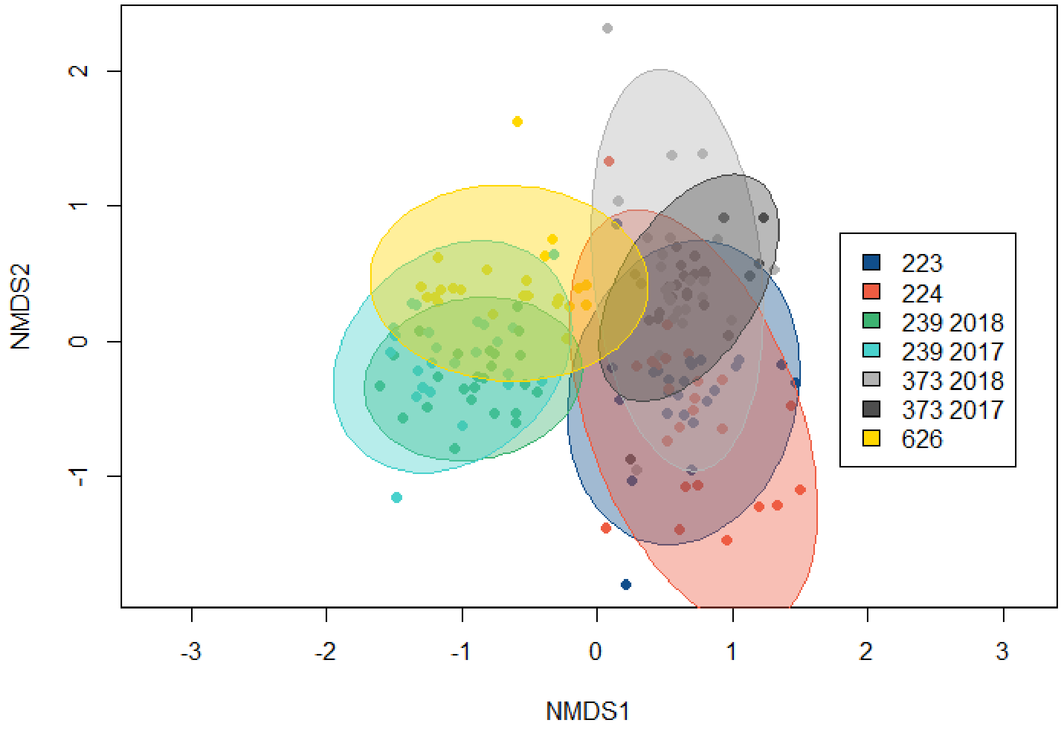

Figure S4: The same nMDS is visualised with 95% confidence ellipses superimposed over the different sample seasons. Samples collected in October were more similar to each other than samples collected in August (as indicated by the smaller ellipses). This is possibly due to greater water mixing taking place in October during autumn turnover which could make the distribution of eNAs more homogeneous within the water samples (see Littlefair, Hrenchuk, Blanchfield, Rennie, & Cristescu, 2021).

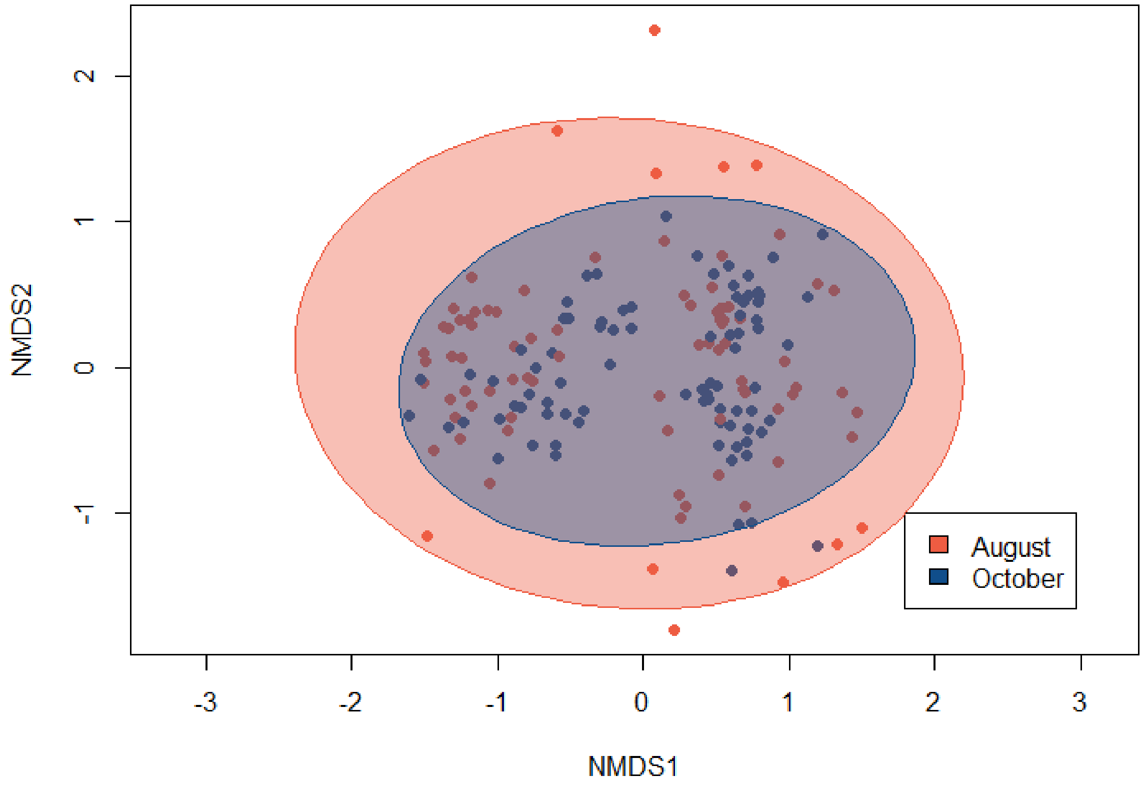

Figure S5: The same nMDS is visualised with 95% confidence ellipses superimposed to group together samples belonging to eDNA and eRNA. Within the entire dataset, eDNA and eRNA detected similar community compositions, although within each lake the two molecules had slight differences (Figure 3 in the main manuscript).

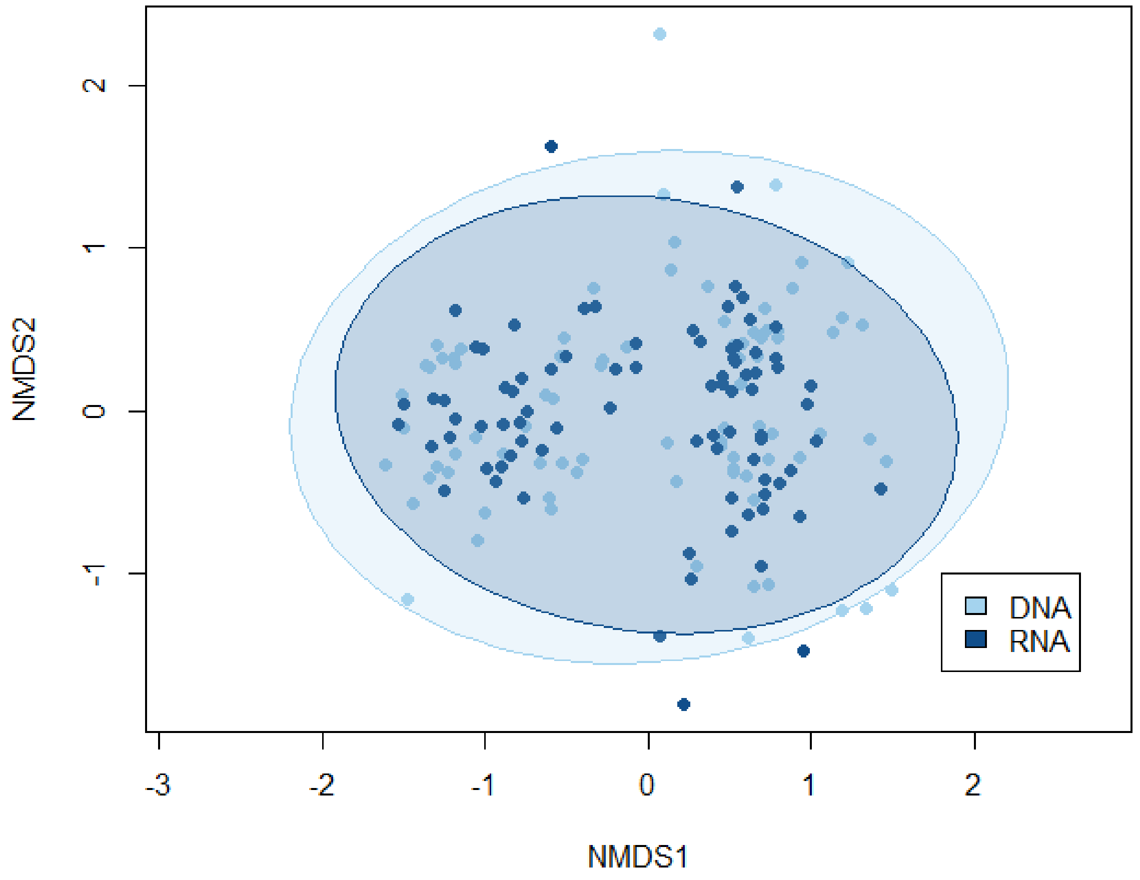
